# Motor planning brings human primary somatosensory cortex into action-specific preparatory states

**DOI:** 10.1101/2020.12.17.423254

**Authors:** Giacomo Ariani, J. Andrew Pruszynski, Jörn Diedrichsen

## Abstract

Motor planning plays a critical role in producing fast and accurate movement. Yet, the neural processes that occur in human primary motor and somatosensory cortex during planning, and how they relate to those during movement execution, remain poorly understood. Here we used 7T functional magnetic resonance imaging (fMRI) and a delayed movement paradigm to study single finger movement planning and execution. The inclusion of no-go trials and variable delays allowed us to separate what are typically overlapping planning and execution brain responses. Although our univariate results show widespread deactivation during finger planning, multivariate pattern analysis revealed finger-specific activity patterns in contralateral primary somatosensory cortex (S1), which predicted the planned finger action. Surprisingly, these activity patterns were as informative as those found in contralateral primary motor cortex (M1). Control analyses ruled out the possibility that the detected information was an artifact of subthreshold movements during the preparatory delay. Furthermore, we observed that finger-specific activity patterns during planning were highly correlated to those during execution. These findings reveal that motor planning activates the specific S1 and M1 circuits that are engaged during the execution of a finger press, while activity in both regions is overall suppressed. We propose that preparatory states in S1 may improve movement control through changes in sensory processing or via direct influence of spinal motor neurons.

**Significance statement:** Motor planning is important for good behavioral performance, yet it is unclear which neural processes underlie the preparation of the nervous system for an upcoming movement. Using high-resolution functional neuroimaging, we investigated how motor planning for finger presses changes the activity state in primary motor and primary somatosensory cortex, and how brain responses during planning and execution relate to each other. We show that planning leads to finger-specific activation in both M1 and S1, which is highly similar to the finger-specific activity patterns elicited during execution. Our findings suggest that S1 is being specifically prepared for an upcoming action, either to actively contribute to the outflowing motor command or to enable action-specific sensory gating.

## Introduction

Animals are capable of generating a wide variety of dexterous behaviors accurately and effortlessly on a daily basis. This remarkable ability relies on the motor system reaching the appropriate preparatory state before each movement is initiated.

At the level of behavior, the process of motor programming, or planning, has long been shown to be beneficial to performance (Keele, 1968; Keele and Summers, 1976; Rosenbaum, 1980), leading to faster reaction times (Klapp and Erwin, 1976; Klapp, 1995; Haith et al., 2016) and more accurate response selection (Ghez et al., 1997; Wong and Haith, 2017; Ariani and Diedrichsen, 2019; Hardwick et al., 2019). The behavioral study of motor planning led to neurophysiological investigations showing the presence of preparatory signals in the patterns of neuronal firing in the dorsal premotor cortex, PMd (Cisek and Kalaska, 2004, 2010; Hoshi and Tanji, 2006), the supplementary motor area, SMA (Hoshi and Tanji, 2004), and the posterior parietal cortex, PPC (Cui and Andersen, 2007, 2011; Andersen and Cui, 2009). Building on this work, human neuroimaging studies have shown that activity in parieto-frontal brain regions during planning of prehension movements can be used to decode several movement properties such as grip type (Gallivan et al., 2011b; Ariani et al., 2015), action order (Gallivan et al., 2015), and effector (Gallivan et al., 2011a, 2013; Leoné et al., 2014; Turella et al., 2016).

At the level of neural population dynamics (Vyas et al., 2020), motor planning can be understood as bringing the neuronal state towards an ideal preparatory point. Once this state is reached and the execution is triggered, the intrinsic dynamics of the system then let the movement unfold (Churchland et al., 2010; Shenoy et al., 2013). Preparatory neural processes have not only been observed in premotor and parietal areas, but also in primary motor cortex (M1, Tanji and Evarts, 1976; Crammond and Kalaska, 2000; Ariani et al., 2018). In contrast, the degree to which primary somatosensory cortex (S1) receives information about the planned movement before movement onset is less clear.

S1 is often considered to be mostly concerned with processing incoming sensory information from tactile and proprioceptive receptors arising after movement onset. Consistent with this notion, previous fMRI studies have not detected the presence of planning-related information in this area (Gallivan et al., 2011a, 2011b, 2013, 2015; although see Gale et al., 2021). However, challenging this notion, in the past years research has shown that S1 can be somatotopically activated even in the absence of tactile inputs, for instance during touch observation (Kuehn et al., 2014), attempted movements without afferent tactile inputs (Kikkert et al., 2021), and through attentional shifts (Puckett et al., 2017). Moreover, a recent human electrocorticography (ECoG) study suggested a possible role for S1 in cognitive-motor imagery (Jafari et al., 2020). The authors recorded neural activity from S1 while a tetraplegic participant imagined reaching movements and found that S1 neurons encoded movement direction during motor imagery in the absence of actual sensations. Another recent ECoG study in non-human primates (Umeda et al., 2019) showed grasp-specific information in the signals from S1 well before movement initiation, and only slightly later than in M1.

However, it remains unknown whether S1 plays a role during motor planning in human participants with an intact sensory system. Furthermore, we currently do not know how the signals during action preparation relate to those during execution, a fact that could provide important insight into the role these signals may play.

Here we designed a high-field (7T) fMRI experiment to study what brain regions underlie the planning of individual finger presses and how these brain representations relate to those during execution. We used variable delays between an instructing cue and a go signal, as well randomly interspersed no-go trials, to temporally separate the evoked responses to movement planning and execution. Using advanced multivariate pattern analyses we were able to examine the relationship between the fMRI patterns related to planned and executed finger actions.

## Results

### Deactivation in sensorimotor regions during planning of finger actions

We instructed 22 participants to plan and execute repeated keypresses with individual fingers of their right hand on a keyboard device while being scanned with 7T fMRI. The key to be pressed corresponded to one of three fingers and was cued during the preparation phase by numbers (1 = thumb, 3 = middle, 5 = little, e.g., Fig. 1A) presented on a computer screen that was visible to the participants lying in the scanner through an angled mirror. After a variable delay (4-8 seconds), participants received a color cue indicating whether to press the planned finger (go trials), or whether to withhold the response (no-go trials). Upon the go cue, participants had to initiate the correct response as fast as possible and make 6 presses of the designated finger, before receiving accuracy points for reward (see Methods).

**Figure 1.**
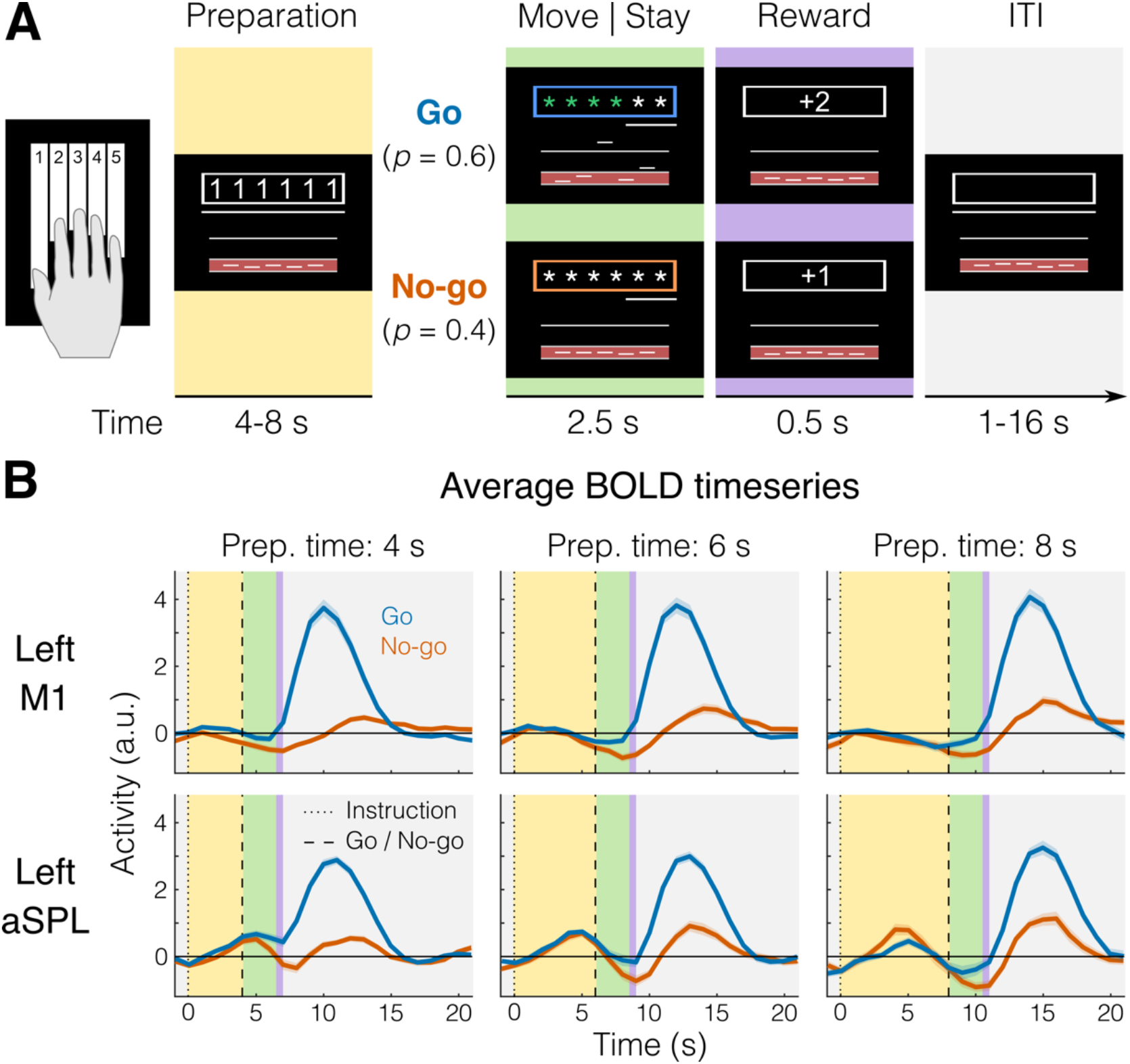
fMRI task and BOLD responses. **A.** Example trial with timing information. Background colors indicate different experimental phases (yellow = preparation; green = move (go) or stay (no-go); purple = reward; gray = inter-trial interval, ITI). **B.** Group-averaged BOLD response (N = 22) for go (blue) and no-go (orange) trials in a region that shows no planning evoked activity (Left M1, top), and one that shows some planning evoked activity (Left aSPL, bottom). Shaded areas indicate standard error of the mean (SEM). Background colors correspond to trial phases as in A.

To control for involuntary overt movements during the preparation phase, we required participants to maintain a steady force on all the keys during the delay, which was closely monitored online. To ensure that planning results would not be biased by the subsequent execution, we restricted all our analyses of the preparation phase to no-go trials only (see Methods). First, we asked which brain regions showed an evoked response during the planning of finger presses (e.g., Fig. 1B).

We focused our analysis on the lateral aspect of the contralateral (left) hemisphere (purple and white areas of the Fig. 2 inset), which included the primary motor and somatosensory cortex, as well as the premotor cortex and anterior parietal cortical regions. To examine brain activation during finger planning, we performed a univariate contrast of the preparation phase (across the three fingers) versus the resting baseline (Fig. 2A). Overall, the instruction stimulus evoked little to no activation in our regions of interest (ROIs, see Methods). In fact, compared to resting baseline, we observed significant deactivation (Fig. 2E) in the primary motor cortex (M1, *t*_21_ = −6.939, *p* = 7.446e-07), the primary somatosensory cortex (S1, *t*_21_ = −5.508, *p* = 1.823e-05), and the dorsal premotor cortex (PMd, *t*_21_ = −2.929, *p* = 0.008). In comparison, execution strongly activated M1 and S1 (Fig. 2C), with activation being significant in all tested ROIs (Fig. 2E, all *t*_21_ > 14.824, all *p* < 1.351e-12).

**Figure 2.**
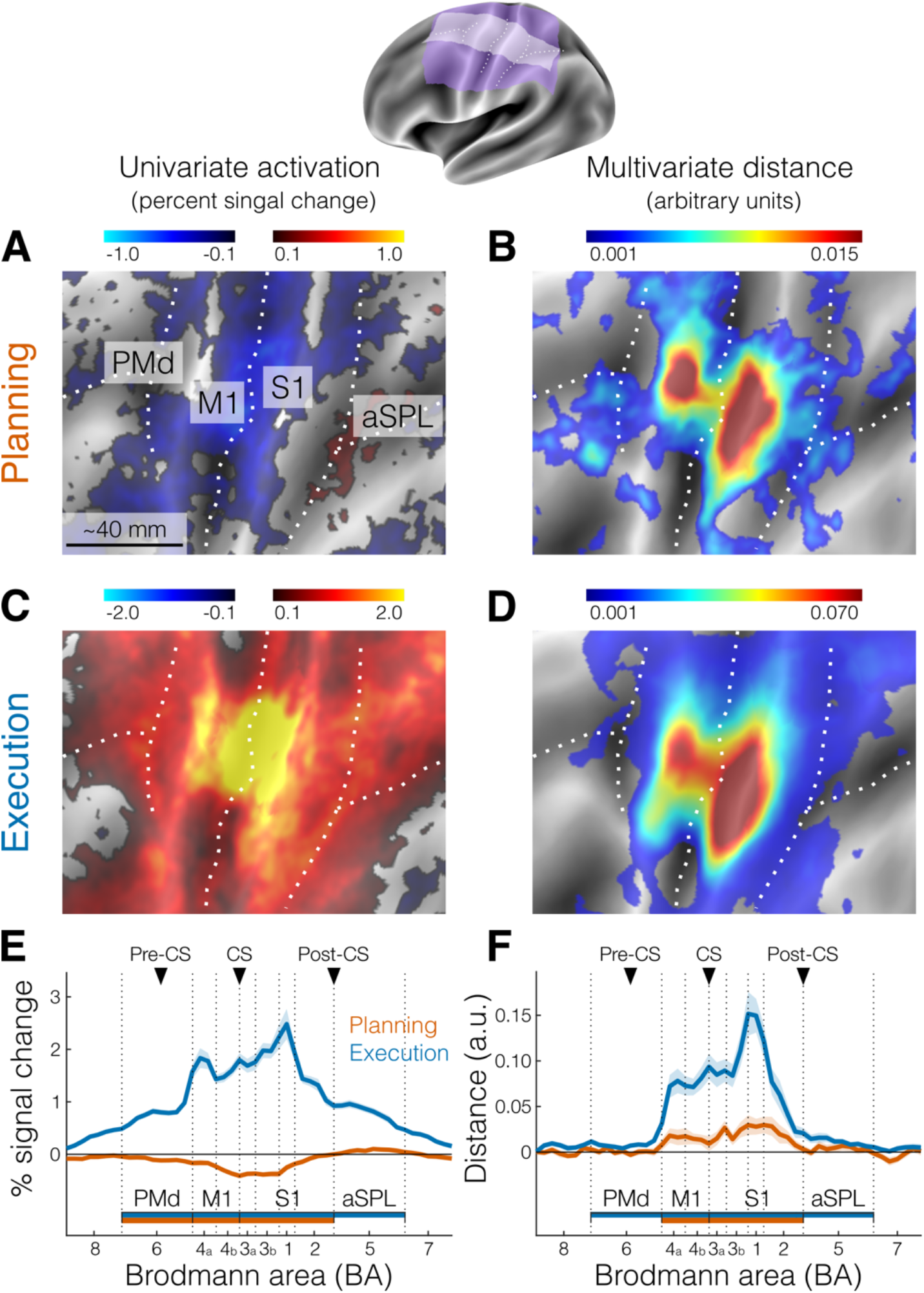
Activation and distance analyses of movement planning and execution. The inset shows the inflated cortical surface of the contralateral (left) hemisphere, highlighting the area of interest (Fig. 2A-B-C-D, purple) and the strip used for the profile ROI analysis (Fig. 2E-F, white). Major sulci are indicated by white dotted lines. **A.** Univariate activation map (percent signal change) for the contrast planning>baseline (no-go trials only). **B.** Multivariate searchlight map of the mean crossnobis distance between the planning of the three fingers (no-go trials only). **C.** Same as A, but for the univariate contrast execution>baseline (go trials). **D.** Same as B, but for the mean crossnobis distance between fingers during execution. Colorbars in A and C reflect mean percent signal change, whereas colorbars in B and D reflect mean crossnobis distance (arbitrary units). **E.** Profile ROI analysis (see Methods) of the mean percent signal change (± SEM) during planning (no-go trials, orange) and execution (blue). The x-axis corresponds to Brodmann areas selected from the white strip shown in the inset at the top. Horizontal bars indicate significance (*p* < 0.05) in a 2-sided one-sample *t*-test vs zero for selected ROIs. **F.** Same as E, but for the mean crossnobis distance (± SEM). Vertical dotted lines mark the approximate boundaries between Brodmann areas subdvisions of our main regions of interest (ROIs, see Methods). Black triangles point to the approximate location of the main anatomical landmarks: Pre-CS = precentral sulcus; CS = central sulcus; Post-CS = postcentral sulcus. PMd (BA 6) = dorsal premotor cortex; M1 (BA 4a, 4b) = primary motor cortex; S1 (BA 3a, 3b, 1, 2) = primary somatosensory cortex; aSPL (BA 5) = anterior superior parietal lobule. For analogous results using the estimates of planning activity from all trials, see Fig. 2 – supplement 1. For the whole brain maps of univariate and multivariate results, see Fig. 2 – supplement 2.

### Planning induces informative patterns in primary somatosensory and motor cortex

Although we found little univariate activation in our main ROIs, preparatory processes need not increase the overall activation in a region. Rather, the region could converge to a specific preparatory neural state (Churchland et al., 2010), while activity increments and decrements within the region (i.e., at a finer spatial scale) average each other out. In this case, information about planned movements would be present in the multivoxel activity patterns in that region. To test this idea, we calculated the crossnobis dissimilarity (or distance, see Methods) between activity patterns. Systematically positive values of this dissimilarity measure indicate that the patterns reliably differentiate between the different planned actions (Walther et al., 2016; Diedrichsen et al., 2020). Indeed, a surface-based searchlight approach (Oosterhof et al., 2012) revealed reliably positive crossnobis distance between the activity patterns related to planning of individual finger presses (Fig. 2B), which the ROI analysis confirmed to be significantly greater than zero in both M1 (*t*_21_ = 2.343, *p* = 0.029) and S1 (*t*_21_ = 3.137, *p* = 0.005, Fig. 2F).

The distribution on the flat surface map of these distance values during planning (Fig. 2B) appeared to be highly similar to the distribution of distances during execution (Fig. 2D). To quantify this topographic similarity, we computed the ratio between distances in different Brodmann area (BA) subdivisions of our ROIs (see Methods), reasoning that a mismatch in location would result in large differences in ratio values. However, the ratio between planning and execution distances was roughly stable across the different subregions of sensorimotor cortex (BA 4a: 0.23, BA 4b: 0.16, BA 3a: 0.24, BA 3b: 0.19, BA 1: 0.22, BA 2: 0.31). In other words, the average distance between finger-specific activity patterns during planning was between 16% and 31% of the average pattern distance during execution. Thus, we not only show the existence of planning related activity in S1, but also that S1 activity patterns are at least as informative as M1 activity patterns.

Visual inspection suggested that the informative patterns during planning may be concentrated more dorsally in M1 and S1 relative to execution. To test for the possibility that the location on the flat surface map of the peaks of the crossnobis distance for M1 and S1 was statistically different across subjects between planning and execution, we used a Hotelling T^2^ test that allowed us to compare the difference between two multivariate means of different distributions (i.e., the distributions of x-y coordinates for the peaks of planning and execution). This test revealed no systematic difference in the location of the peak vertex between planning and execution across subjects (M1: *T^2^*_2,20_ = 0.725, *p* = 0.712; S1: *T^2^*_2,20_ = 2.424, *p* = 0.335).

Together, our analyses indicate that information about single finger actions is already represented during motor planning in the same parts of the primary motor and somatosensory cortices that are engaged during execution of the presses. Given that we only used the activity estimates from no-go trials (~40% of total trials), this information cannot be explained by a spill-over from subsequent execution-related activity. An analysis using the estimates of planning activity from all trials yielded very similar results (see Fig. 2 - supplement 1), demonstrating that we could separate planning from execution-related signals.

### Activity patterns are not caused by small movements during the preparation phase

The presence of planning-related information in primary sensorimotor regions was surprising, especially in S1, where it had not previously been reported in comparable fMRI studies (Gallivan et al., 2011b, 2015). To ensure that these results were not caused by overt movement, participants were instructed to maintain a steady force on the keyboard during the preparation phase, such that we could monitor even the smallest involuntary preparatory movements.

Inspection of the average force profiles (Fig. 3A) revealed that participants were successful in maintaining a stable force between 0.2 and 0.4 N during preparation. However, averaging forces across trials may obscure small, idiosyncratic patterns visible during individual trials (Fig. 3B) that could be used to distinguish the different movements. To test for the presence of such patterns, we submitted both the mean and standard deviation of the force traces on each finger to a multivariate dissimilarity analysis (see Methods). Indeed, this sensitive analysis revealed that some participants showed small movement patterns predictive of the upcoming finger (positive behavioral distances in Fig. 3C).

**Figure 3.**
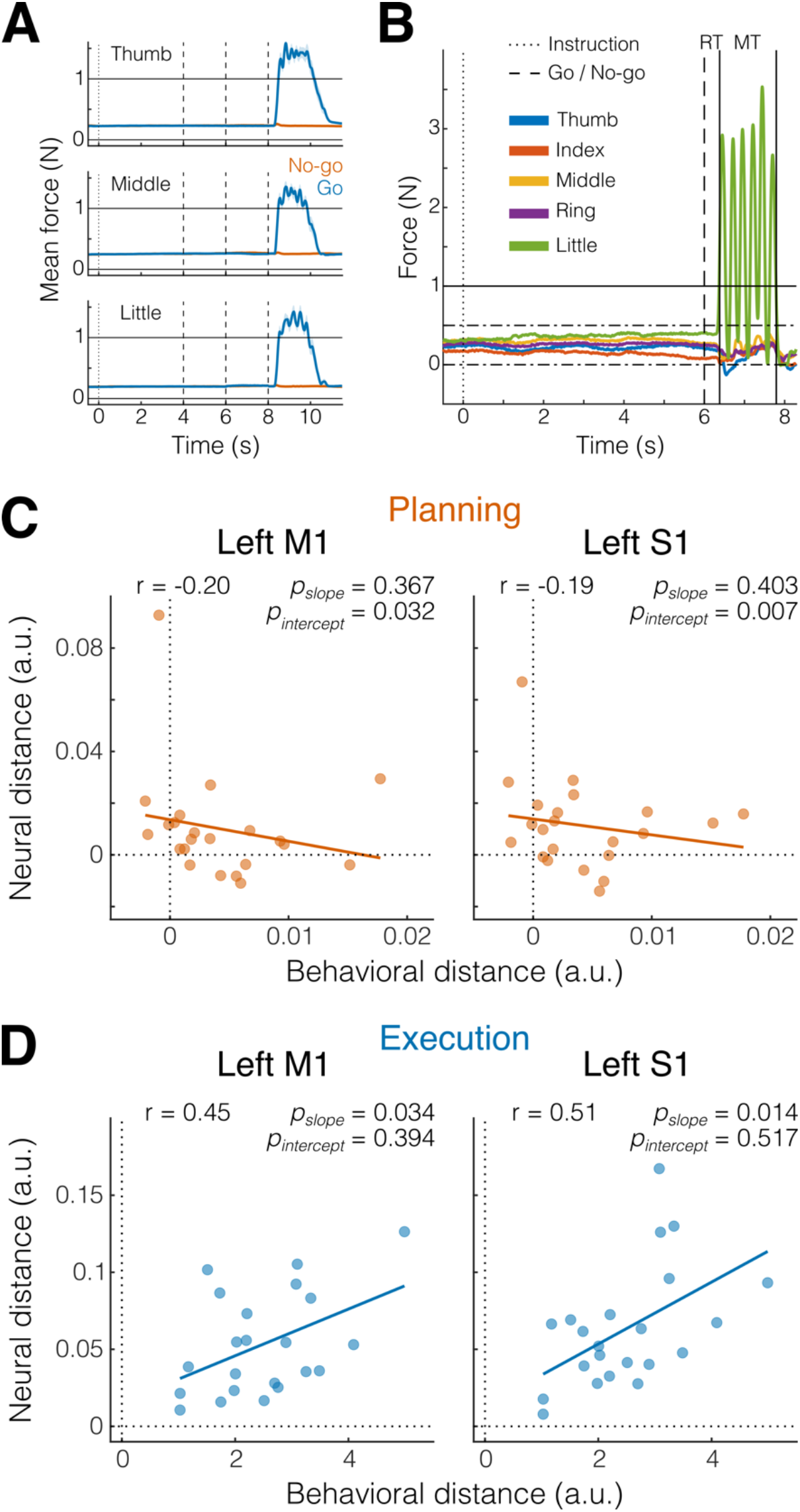
Small involuntary movements do not explain preparatory activity patterns in M1 and S1. **A.** Mean finger force (± SEM) plotted in 10 ms bins, time-aligned to instruction onset (dotted vertical line) and end of the preparation phase (dashed vertical lines), separately for the three fingers and go (blue) and no-go (orange) trials. **B.** Example of an individual trial with a 6 s preparation phase, followed six presses of the little finger (green). Horizontal solid line denotes press threshold (1 N). Dash-dotted lines denote the boundaries of the finger preactivation red area in Fig. 1A (see Methods). Reaction time (RT) was defined as the time from the go cue (dashed vertical line) to the onset of the first press (left solid vertical line). Movement time (MT) was defined as the time from the onset of the first press (left solid vertical line) until the release of the last press (right solid vertical line). **C.** Pearson’s correlation (r) between behavioral and neural distances in M1 and S1 (see Methods) during the preparation phase (planning, orange). Each dot represents an individual participant (N = 22). Solid line shows linear regression fit; *p*-values pairs refers to the slope and the intercept of the fitted line. **D.** Same as C, but during the movement phase (execution, blue).

These distances, however, were ~200-300 times smaller than the average distances during execution (x-axis in Fig. 3D), and we found no significant correlation between the magnitude of the behavioral differences for the preparation phase and the amount of planning information present in our sensory-motor regions of interest (both *p*-values for the slope of the linear fit > 0.3 in Fig. 3C). More importantly, a significantly positive intercept in the linear fit in Fig. 3C (M1: *p* = 0.032; S1: *p* = 0.007) shows that, even after correcting for the influence behavioral patterns, the activity patterns in M1 and S1 remained informative (i.e., significantly positive neural distance even with no significant behavioral distance). Thus, the finding of finger-specific activity patterns in M1 and S1 cannot be explained by small involuntary movements during the preparation phase.

### Single finger activity patterns from planning to execution are positively correlated

How do planning-related activity patterns in M1 and S1 relate to the activity patterns observed during execution? Neurophysiological experiments have suggested that patterns of movement preparation are orthogonal – or uncorrelated – to the patterns underlying active movement (Kaufman et al., 2014). This arrangement allows movement preparation to occur without causing overt movement.

When we compared the planning- and execution-related activity patterns as measured with fMRI, a technique that samples neuronal activity at a much coarser spatial resolution, we found the opposite result. Planning- and execution-related patterns for the same finger were tightly related.

First, inspection of the representational dissimilarity matrices (RDMs) for M1 and S1 (Fig. 4A) shows a large difference between planning and execution patterns, which is due to the substantially higher average activation during movement compared to planning. This overall distance between planning and execution can also be appreciated in a 3D projection of the RDMs using multidimensional scaling (MDS) to highlight the representational geometry between activity patterns (principal component PC1 in Fig. 4B, *top*).

**Figure 4.**
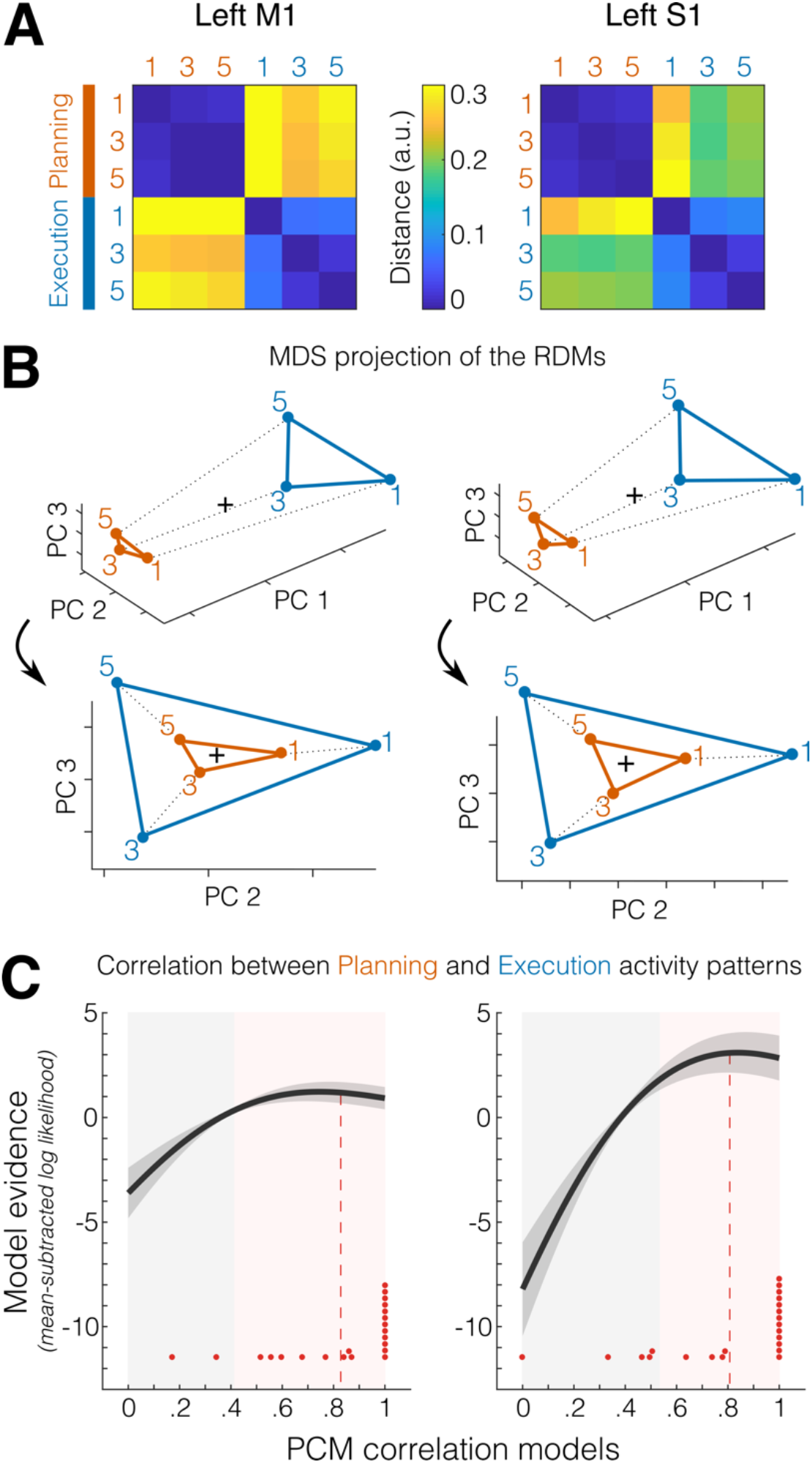
Correlated representations of single fingers across planning and execution. **A.** Representational dissimilarity matrices (RDMs) of the activity patterns evoked by the three fingers during the preparation (no-go planning, orange) and movement (execution, blue) phases, separately for the two main ROIs (Left M1, *left*; Left S1, *right*). **B.** Multidimensional scaling (MDS) projection of the RDMs in A. *Top*, 3D view highlighting the first principal component (PC 1, difference in average activation between planning and execution). *Bottom*, rotated view highlighting the correspondence between representational geometries across planning and execution visible on PC 2 and PC 3. **C.** Pattern component modelling (PCM) evaluation of models (x-axis) of different correlations between planning- and execution-related activity patterns. Shown in dark gray is the group-average of the individual log-likelihood (± SEM across participants) curves expressed as a difference from the mean log-likelihood across models (i.e., zero on the y-axis). Red dots indicate the best fitting correlation model for each participant (N = 22). Red dashed lines denote the average winning (i.e., best fitting) models across participants. Gray-shaded areas indicate models that perform statistically worse (*p* < 0.05) than the best fitting correlation model (determined in a cross-validated fashion, see Methods). Pink-shaded areas indicate the correlation models that perform statistically equivalent (*p* ≥ 0.05) to the best fitting correlation model.

Second, within each phase, the pattern for the thumb was more distinct than those for the other fingers, replicating previous results from execution alone (Ejaz et al., 2015; Yokoi et al., 2018). Importantly, however, when ignoring the overall difference between the mean patterns for planning and execution, by looking at a rotated view of the representational geometry (Fig. 4B, *bottom*), it became clear that the finger patterns were arranged in a congruent way, with planning and execution related activity patterns for the same finger being closer to one another. This representation suggested that the finger-specific patterns during planning may be a scaled-down version of the patterns during execution.

To test this idea more precisely, we quantified the correspondence (i.e., correlation) between planning and execution pattern for each finger using Pattern Component Modeling (PCM, Diedrichsen et al., 2018). Simple correlations between measured fMRI patterns are usually lower than the true correlation because of the biasing influence of measurement noise (see http://www.diedrichsenlab.org/BrainDataScience/noisy_correlation/index.htm). PCM corrects for this effect by evaluating the likelihood of the data (taking into account the measurement noise), assuming true correlation between 0 and 1 (see Methods for details).

From the individual fits (Fig. 4C, red dots), we found that the most likely correlation model across participants was at 0.83 (± 0.053 SEM) for M1 and at 0.81 (± 0.061 SEM) for S1 (Fig. 4C, red dashed lines). By comparing these estimates to the zero-correlation model, we can conclude that the correlation of finger-specific patterns across planning and execution was significantly larger than zero in both M1 and S1 (both *t*_21_ > 13.288, *p* < 1.086e-10 in a two-tailed one sample *t*-test against zero). However, the maximum likelihood estimates of the correlation cannot be used to evaluate whether the overlap of these patterns was only partial (r < 1) or complete (r = 1), as the estimates are biased due to measurement noise (Walther et al., 2016; Diedrichsen et al., 2018).

We therefore compared the log-likelihoods of the best fitting model (computed using a cross-validated approach, see Methods) to the log-likelihoods under the model that the two are perfectly correlated (r = 1). Across participants, this difference was not significant for either M1 (*t*_21_ = 0.953, *p* = 0.176) or S1 (*t*_21_ = 0.148, *p* = 0.442). Given that no correlation model had significantly higher log-likelihoods than the 1-correlation model, we cannot rule out that the underlying true correlation was indeed 1. In other words, we have as much evidence that the correspondence was only partial as we do that the correspondence was perfect. By comparing the best fitting correlation model to every other correlation model, we have evidence that the true (i.e., noiseless) correlation between planning and execution finger-specific activity pattern was between 0.41–1.0 in M1 and between 0.54–1.0 in S1 (Fig. 4C, pink-shaded areas).

Thus, our data is consistent with the idea that, at the resolution of fMRI, the activity patterns for planning and execution of finger presses in S1 and M1 are either partially overlapping or even a scaled version of each other.

## Discussion

In the present study, we asked participants to produce repeated single finger presses while undergoing 7T fMRI. We used variable preparatory delays and no-go trials to cleanly dissociate the brain responses to the consecutive preparation and movement phases. We found that information about planned finger actions is present in both S1 and M1 before action onset, even though the overall level of activation in these regions was below resting baseline. Moreover, while execution elicited much higher brain activation, the fine-grained, finger-specific activity patterns were highly similar across planning and execution. Control analyses confirmed that the observed results were not caused by pre-movement finger activity.

Our finding that motor planning activates M1 in a finger-specific fashion was not necessarily surprising given many neurophysiological studies reporting anticipatory activity of M1 neurons related to movement intentions (Tanji and Evarts, 1976; Riehle and Requin, 1989; Alexander and Crutcher, 1990), as well as human neuroimaging showing shared information between delayed and immediate movement plans (Ariani et al., 2018). In contrast, the robust activity patterns related to single finger planning in S1 were more surprising, given that this region has classically been associated with the passive processing of somatosensory information from receptors in the skin, muscles, and tendons.

So, what could then be the role of S1 during movement planning? First, it is worth noting that there are substantial projections from S1 (Brodmann area 3a) that terminate in the ventral horn of the cortico-spinal tract (Coulter and Jones, 1977; Rathelot and Strick, 2006). Although stimulation of area 3a in macaques typically fails to evoke overt movements (Widener and Cheney, 1997), it has been suggested that this population of cortico-motoneurons specifically projects to gamma motoneurons that control the sensitivity of muscle spindle afferents (Rathelot and Strick, 2006). Thus, it is possible that S1 plays an active role in movement generation by preparing the spindle apparatus in advance of the movement.

Second, the finger-specific preparatory state in S1 may reflect the prediction of the upcoming sensory stimulation, allowing for a movement-specific sensory gain control (Azim and Seki, 2019). It is likely that, this process is also accompanied by an allocation of attention to the cued finger. However, as voluntary planning requires attention, our current dataset cannot distinguish between the two possibilities. Sensory stimuli could become attenuated to maintain movement stability and filter out irrelevant or self-generated signals. Indeed, multiple studies have shown that both somatosensation and somatosensory-evoked potentials in S1 decrease during voluntary movement (Starr and Cohen, 1985; Chapman et al., 1987; Jiang et al., 1990; Seki and Fetz, 2012). Alternatively, sensory processing of the expected salient signals could be enhanced to improve movement execution.

While several previous fMRI studies did not find action-specific encoding in S1 during planning (Gallivan et al., 2011a, 2011b, 2013, 2015), concurrently with our study a second paper found movement-specific modulation of S1 preparatory activity (Gale et al., 2021). Together, these two papers provide convergent evidence that motor planning triggers notable changes in the neural state of the somatosensory system and that such changes can be detected with fMRI in humans.

The second important finding in our paper was the close correspondence between finger-specific activity patterns across planning and execution—which appears to be at odds with the idea that these two processes occupy orthogonal neural subspaces to avoid overt movement during planning (Kaufman et al., 2014; Elsayed et al., 2016). We think that there are at least two possible explanations for this. First, the divergence of results could be caused by the difference in behavioral paradigms. While the neuronal correlates of movement planning in non-human primates have largely been studied using upper limb movements, we used here individuated finger presses. If for single finger actions even single-neuron activity patterns are highly correlated between planning and execution, then overt movement during planning would need to be actively suppressed, for example through the deactivation that we observed around the central sulcus.

An alternative and perhaps more likely explanation of the discrepancy lies in the different measurement modalities. Orthogonality was observed in electrophysiological recordings of individual neurons, whereas the fMRI measurements we employed here mainly reflect excitatory postsynaptic potentials (Logothetis et al., 2001) and average metabolic activity across hundreds of thousands of cortical neurons. Thus, it is possible that planning pre-activates the specific cortical columns in M1 and S1 that are also most active during movement of that finger. Within each of these cortical micro-circuits, however, planning-related activity could still be orthogonal to the activity observed during execution at the single neuron level (e.g., see Arbuckle et al., 2020, for a similar observation for cortical representations of flexion and extension finger movements). This would suggest a new hypothesis for the architecture of the sensory-motor system where movement planning pre-activates the action-specific circuits in M1 and S1. However, it does so in a fashion that the induced planning-related activity is, in terms of the firing output of neurons, orthogonal to the patterns during execution.

## Methods

### Participants

Twenty-three right-handed neurologically healthy participants volunteered to take part in the experiment (13 F, 10 M; age 20–31 years, mean 23.43 years, SD 4.08 years). Criteria for inclusion were right-handedness and no prior history of psychiatric or neurological disorders. Handedness was assessed with the Edinburgh Handedness Inventory (mean 82.83, SD 9.75). All experimental procedures were approved by the Research Ethics Committee at Western University. Participants provided written informed consent to procedures and data usage and received monetary compensation for their participation. One participant withdrew before study completion and was excluded from data analysis (final N = 22).

### Apparatus

Repeated right-hand finger presses were performed on a custom-made MRI-compatible keyboard device (Fig. 1A). Participants only used the tips of their fingers to press on the keys. The keys of the device did not move but force transducers underneath each key measured isometric force production at an update rate of 2 ms (Honeywell FS series; dynamic range 0-25 N; sampling 200 Hz). A keypress/release was detected when the force crossed a threshold of 1 N. The forces measured from the keyboard were low pass filtered to reduce noise induced by the MRI environment, amplified, and sent to PC for online task control and data recording.

### Task

We used a task in which participants produced repeated keypresses with the tip of their right-hand fingers in response to numerical cues appearing on a computer screen (white outline, Fig. 1A). On each trial, a string of 6 numbers (instructing cue) instructed which finger press to plan (1 = thumb, 3 = middle, 5 = little).

The length of the preparation phase (yellow background in Fig. 1) was randomly sampled to be 4 s (56% of trials), 6 s (30%), or 8 s (14%). To limit and monitor unwanted movements during the preparation phase, we required participants to pre-activate their fingers by maintaining a steady force of around 0.2-0.3 N on all of the keys during the preparation phase. As a visual aid, we displayed a red area (between 0 and 0.5 N) and asked participants to remain in the middle of that range with all the fingers (touching either boundary of the red area would count as unwanted movement, thus incurring an error).

At the onset of the movement phase (green background), participants received a color cue (go/no-go cue) indicating whether to perform the planned finger presses (blue outline = go, *p* = 0.6), or not (orange outline = no-go, *p* = 0.4). The role of no-go trials was to de-couple the hemodynamic response to the successive planning and execution events, which would otherwise always overlap in fast fMRI designs due to the sluggishness of the fMRI response (Ariani et al., 2018).

To encourage planning during the delay period, at the go cue the digits were masked with asterisks, and participants had to perform the presses from memory. Participants had 2.5 s to complete the movement phase, and a vanishing white bar under the asterisks indicated how much time was left to complete all of the keypresses. Participants received online feedback on the correctness of each press with asterisks turning either green, for a correct press, or red, for incorrect presses. As long as the participants remained within task constraints (i.e., 6 keypresses in less than 2.5 s), an exact movement speed was not enforced. In no-go trials, participants were instructed to remain as still as possible maintaining the finger pre-activation until the end of the movement phase (i.e., releasing any of the keys would incur an error).

During the reward phase (0.5 s, purple background) points were awarded based on performance and according to the following scheme: −1 point in case of no-go error or go cue anticipation (timing errors); 0 points for pressing any wrong key (press error); 1 point in case of a correct no-go trial; and 2 points in case of a correct go trial.

Inter-trial-intervals (ITI, gray background) were randomly drawn from {1, 2, 4, 8, 16 s} with the respective proportion of trials {0.52, 0.26, 0.13, 0.6, 0.3}.

### Experiment design and structure

Our chosen distribution of preparation times, inter-trial intervals, and no-go trials, was determined by minimizing the variance inflation factor (VIF) for a given length of scan:

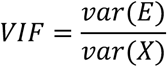

Where ***var(E)*** is the mean estimation variance of all the regression weights (planning- and execution-related regressors for each finger), and ***var(X)*** the mean estimation variance had these regressors been estimated in isolation. The VIF quantifies the severity of multicollinearity between model regressors by providing an index of how much the variance of an estimated regression coefficient is increased because of collinearity. Large values for VIF mean that model regressors are not independent of each other, whereas a VIF of 1 means no inflation of variance. After optimizing the design, the VIF was quite low, on average around 1.15, indicating that we could separate planning and execution related activity without a large loss of experimental power.

Participants underwent one fMRI session consisting of 10 functional runs and 1 anatomical scan. In an event-related design, we randomly interleaved 3 types of repeated single finger presses involving the tip of the thumb (1), the middle (3), and the little (5) fingers (e.g., 111111 for thumb presses, Fig. 1A) and 3 types of multi finger sequences (e.g., 135315).

The day before the fMRI scan, participants familiarized themselves with the experimental apparatus and the go/no-go paradigm in a short behavioral session of practice outside the scanner (5 blocks, about 15-30 min in total). For the behavioral practice, inter-trial intervals were kept to a fixed 1 s to speed up the task, and participants were presented with different sequences from what they would see while in the scanner. These 6-item sequences were randomly selected from a pool of all possible permutations of the numbers 1, 3, and 5, with the exclusion of sequences that contained consecutive repetitions of the same number. Given that the current paper is concerned with the relationship between representations of simple planning and execution, here we will focus only on the results for single finger actions. The results for multi finger sequences are intended for publication in a future paper.

Each single finger trial type (e.g., 111111) was repeated 5 times (2 no-go and 3 go trials), totalling 30 trials per functional run. Two periods of 10 s rests were added at the beginning and at the end of each functional run to allow for signal relaxation and provide a better estimate of baseline activation. Each of the 10 functional runs took about 5.5 minutes and the entire scanning session (including the anatomical scan and setup time) lasted for about 75 minutes.

### Imaging data acquisition

High-field functional magnetic resonance imaging (fMRI) data were acquired on a 7T Siemens Magnetom scanner with a 32-channel head coil at Western University (London Ontario, Canada). The anatomical T1-weighted scan of each participant was acquired halfway through the scanning session (after the first 5 functional runs) using a Magnetization-Prepared Rapid Gradient Echo sequence (MPRAGE) with voxel size of 0.75×0.75×0.75 mm isotropic (field of view = 208 x 157 x 110 mm [A-P; R-L; F-H], encoding direction coronal). To measure the blood-oxygen-level dependent (BOLD) responses in human participants, each functional scan (330 volumes) used the following sequence parameters: GRAPPA 3, multi-band acceleration factor 2, repetition time [TR] = 1.0 s, echo time [TE] = 20 ms, flip angle [FA] = 30 deg, slice number: 44, voxel size: 2×2×2 mm isotropic. To estimate and correct for magnetic field inhomogeneities, we also acquired a gradient echo field map with the following parameters: transversal orientation, field of view: 210 x 210 x 160 mm, 64 slices, 2.5 mm thickness, TR = 475 ms, TE = 4.08 ms, FA = 35 deg.

### Preprocessing and univariate analysis

Preprocessing of the functional data was performed using SPM12 (fil.ion.ucl.ac.uk/spm) and custom MATLAB code. This included correction for geometric distortions using the gradient echo field map (Hutton et al., 2002), and motion realignment to the first scan in the first run (3 translations: x, y, z; 3 rotations: pitch, roll yaw). Due to the short TR, no slice timing corrections were applied. The functional data were co-registered to the anatomical scan, but no normalization to a standard template or smoothing was applied. To allow magnetization to reach equilibrium, the first four volumes of each functional run were discarded. The pre-processed images were analyzed with a general linear model (GLM). We defined separate regressors for each combination of the 6 finger-actions (single, multi) x 2 phases (preparation, movement). To control for the effect of potential overlap between execution activity and the preceding planning activity, we also estimated a separate GLM with separate regressors for the preparation phases of go and no-go trials, resulting in a total of 18 regressors (12 go + 6 no-go), plus the intercept, for each run. Each regressor consisted of a boxcar function (on for 2 s of each phase duration and off otherwise) convolved with a two-gamma canonical hemodynamic response function with a peak onset at 5 s and a post-stimulus undershoot minimum at 10 s (Fig. 1B).

Given the relatively low error rates (i.e., number of error trials over total number of trials, timing errors: 7.58 ± 0. 62 %; press errors: 1.18 ± 0.26 %, see Task above), all trials were included to estimate the regression coefficients, regardless of whether the execution was correct or erroneous. Ultimately, the first-level analysis resulted in activation images (beta maps) for each of the 18 conditions per run, for each of the participants.

### Surface reconstruction and ROI definition

Individual subject’s cortical surfaces were reconstructed using Freesurfer (Dale et al., 1999). First, we extracted the white-gray matter and pial surfaces from each participant’s anatomical image. Next, we inflated each surface into a sphere and aligned it using sulcal depth and curvature information to the Freesurfer average atlas (Fischl et al., 1999). Both hemispheres in each participant were then resampled into Workbench’s 164k vertex grid. This allowed us to compare similar areas of the cortical surface in each participant by selecting the corresponding vertices on the group atlas.

Anatomical regions of interest (ROIs) were defined using a probabilistic cytoarchitectonic atlas (Fischl et al., 2008) projected onto the common group surface. Our main ROIs were defined bilaterally as follows: primary motor cortex (M1) was defined by including nodes with the highest probability of belonging to Brodmann areas (BA) 4a and 4b, within 2 cm above and below the hand knob anatomical landmark (Yousry, 1997); primary somatosensory cortex (S1) was defined by the nodes related to BA 1, 2, 3a, and 3b; dorsal premotor cortex (PMd) was defined at the junction between the superior frontal sulcus and the precentral sulcus (BA 6); finally, the anterior part of the superior parietal lobule (aSPL, BA 5) included areas anterior, superior and ventral to the intraparietal sulcus (IPS). ROI definition was carried out separately in each subject using FSL’s subcortical segmentation. When resampling functional onto the surface, to avoid contamination between M1 and S1 activities, we excluded voxels with more than 25% of their volume in the grey matter on the opposite side of the central sulcus.

### Multivariate distance analysis

To detect single finger representations across the cortical surface, we used representational similarity analysis (RSA; Diedrichsen and Kriegeskorte, 2017; Walther et al., 2016) with a surface-based searchlight approach (Oosterhof et al., 2011). For each node, we selected a region (the searchlight) corresponding to 100 voxels (12 mm disc radius) in the gray matter and computed cross-validated Mahalanobis (crossnobis, Walther et al., 2016) dissimilarities between pairs of evoked activity patterns (beta estimates from first level GLM) of single finger sequences, during both preparation and movement phases.

Prior to calculating the dissimilarities, beta weights for each condition were spatially pre-whitened (i.e., weighted by the matrix square root of the noise covariance matrix, as estimated from the residuals of the GLM. The noise covariance matrix was slightly regularized towards a diagonal matrix (Ledoit and Wolf, 2004). Multivariate pre-whitening has been shown to increase the reliability of dissimilarity estimates (Walther et al., 2016).

The resulting analyses (one RDM per participant containing the dissimilarities between the three single fingers during planning and execution: 6 conditions, 15 dissimilarity pairs) were then assigned to the central node and the searchlight was moved across all nodes across the surface sheet obtaining a cortical map (Fig. 2B-2D). Cross-validation ensures the distances estimates are unbiased, such that if two patterns differ only by measurement noise, the mean of the estimated value would be zero. This also means that estimates can sometimes become negative. Therefore, dissimilarities significantly larger than zero indicate that two patterns are reliably distinct, similar to an above-chance performance in a cross-validated pattern-classification analysis.

The searchlight analysis was mainly used for visualization purposes. Additionally, we conducted the multivariate analysis separately for each anatomically defined ROI (e.g., Fig. 4A). For the profile ROI analysis (both univariate and multivariate, e.g., Fig. 2E-F), we defined 50 rectangular surface-based searchlights in each hemisphere that covered the virtual strip shown in the top inset of Fig. 2 and that were aligned to the boundaries between different ROI subdivisions. Based on these surface-based searchlights, we defined the voxel-based subdivisions in individual brains. For statistical comparisons, these subdivisions were successively grouped by averaging within-ROI subdivisions (see Fig. 2E-F). This approach allowed us to compute both ROI-level statistical comparisons and the analysis of the ratio of distances in the different subdivisions of our main ROIs (e.g., M1 into BA4a and BA4b). Statistical comparisons consisted of 2-sided one-sample *t*-test vs zero for selected ROIs.

### Correlation between behavioral and neural distances

To ensure that our planning results were not contaminated by unwanted micro-movements during the preparation phase, we calculated the behavioral distance between the different fingers on the basis of keyboard force data and correlated behavioral and neural distances.

For behavioral distances, we first extracted force data (2 ms temporal resolution, smoothed with a gaussian kernel of 9.42 full width at half maximum, FWHM) and binned it in 10 ms steps (down sampling largely due to memory constraints) for both the preparation and movement phases (Fig. 3A). Next, for each subject, we calculated the mean (5) and the standard deviation (5) of the time-averaged force of each finger for each condition (3 sequences x 2 phases = 6) and block (10). These subject-specific finger force patterns (60 x 10) were multivariately pre-whitened using their covariance matrix. Finally, we calculated the cross-validated squared Euclidean distances for each condition (6 x 6 RDM) and averaged distances between the 3 finger presses for each phase (preparation, movement).

These mean finger force distances for each subject were correlated with the mean voxel activity distances from the two phases for 2 ROIs (M1 and S1, Fig. 3C-3D). To statistically assess that the neural distances were still significantly larger than zero even in the absence of behavioral distances, we computed the *p*-value for the intercept of the linear fit.

### Pattern component modelling correlation models

Visual inspection of the multidimensional scaling (MDS) plots (Fig. 4B) suggested that the finger-specific activity patterns during planning and execution might be arranged in a congruent fashion. To systematically quantify the correspondence of finger-specific activity patterns across planning and execution, we used pattern component modelling (PCM).

This method has been shown to be advantageous in estimating correlations (Diedrichsen et al., 2018). In contrast to simple Pearson’s or cross-validated correlation estimated from raw activity patterns, PCM separately models the noise and signal components.

We created 100 correlation models with correlations in the range [0–1] in equal step sizes and assessed the likelihood of the observed data (Y, the 6 activation patterns observed in 10 runs) from each participant under each correlation (r): *p*(*Y|r*). Apart from a fixed correlation, each model contained 5 parameters, each describing the variance of specific pattern component. The first two parameters captured the variance of the common pattern for all execution patterns and the variance of the common pattern for all planning patterns. Together, these two components captured the overall difference between planning and execution. The next two parameters captured the variance associated with the 3 fingers under the two conditions. Finally, the noise parameter determined the variance of the measurement noise. Because all correlation models had the same number of parameters, we simply maximized the likelihood for each correlation model in respect to these parameters. A full example of a PCM correlation model can be found online at https://pcm-toolbox-python.readthedocs.io/en/latest/demos/demo_correlation.html.

The curve in Fig. 4C shows the average log-likelihood for each correlation model (100 models from 0 to 1 in equal steps sizes), relative to the mean log-likelihood across models (zero on the y-axis). Differences between the log-likelihoods can be interpreted as log-Bayes factors. Group inferences were performed using a simple *t*-tests on the log-likelihoods.

To compare specific models to the best fitting model, we had to correct for the bias arising from picking the best model and testing it on the same data. Therefore we used N-1 subjects to determine the group winning model, and then chose the log-likelihood of this model for the left-out subject (for whom this model may not be the best one) as the likelihood for the “best” model. This was repeated across all subjects and a one-sided paired-sample *t*-test was performed on the recorded log-likelihood and every other model.

This test revealed which of the correlation models were significantly worse (i.e., associated with a lower log-likelihood) than the winning model that was independently estimated via cross-validation (gray-shaded area in Fig. 4C).

## Author contributions

G.A. and J.D. designed research; G.A. performed research; G.A. analyzed data; G.A. drafted the manuscript; G.A., J.A.P, and J.D. edited the manuscript.

## Acknowledgements

This work was supported by a NSERC Discovery Grant (RGPIN-2016-04890) awarded to J.D., and the Canada First Research Excellence Fund (BrainsCAN). The authors wish to thank Eva Berlot for helpful discussions and contributions to data analysis.

## Supplementary figures

**Figure 2 – supplement 1.**
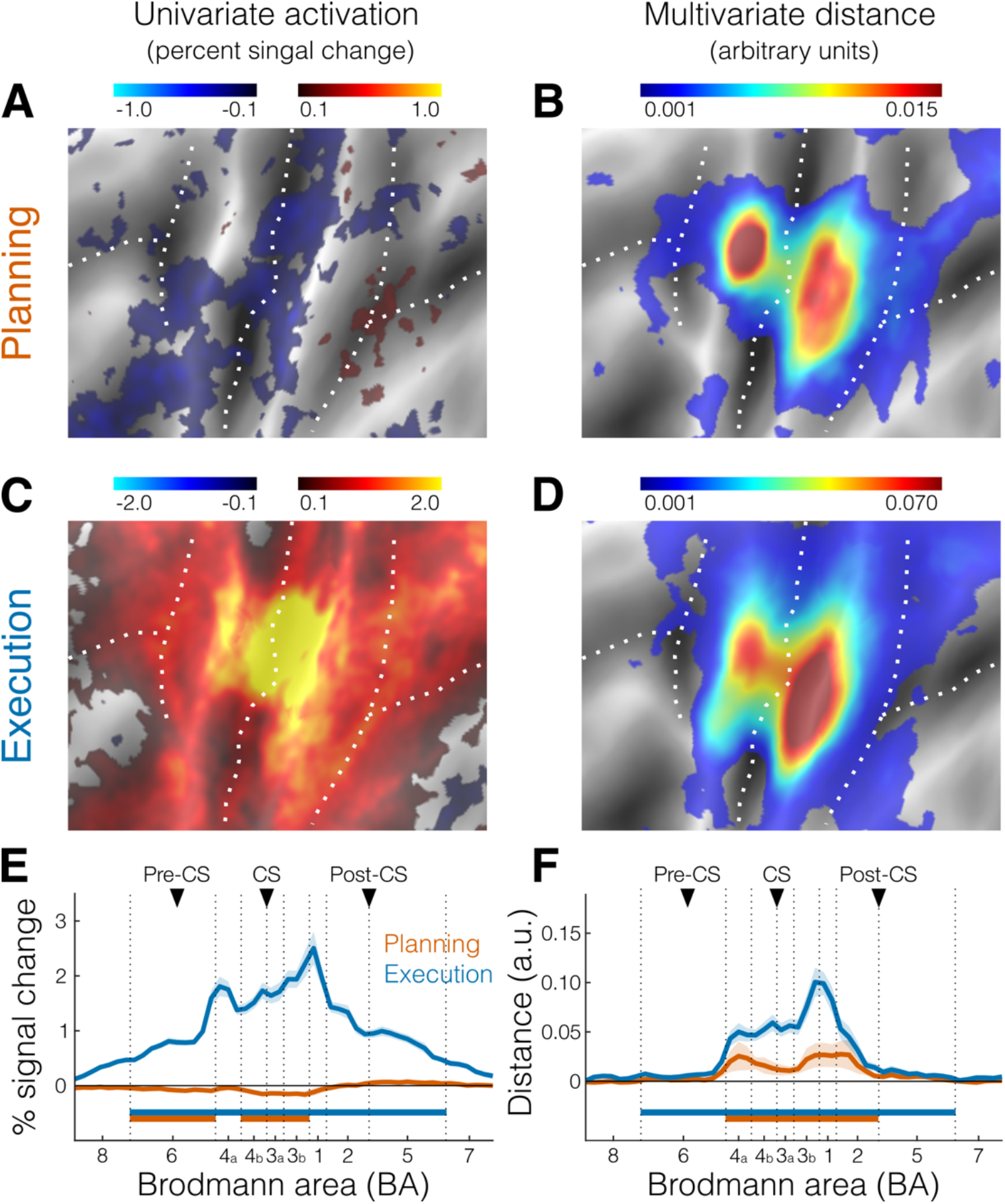
Activation and distance analyses using planning of both go and no-go trials. **A.** Activation map (percent signal change) for the contrast planning>baseline. The selected area of interest is the same as shown in purple in the inset of Fig. 2A. **B.** Crossnobis distance searchlight map for movement planning. **C.** Same as A, but for the contrast execution>baseline. **D.** Same as B, but for movement execution. **E-F.** Profile ROI analysis corresponding to the same area shown in white in the inset of Fig. 2A. **E.** Mean percent signal change (± SEM) during planning (orange) and execution (blue). **F.** mean crossnobis distance (± SEM). Horizontal bars indicate significance (*p* < 0.05) in a 2-sided one-sample *t*-test against zero. All other figure conventions are the same as in Fig. 2.

**Figure 2 – supplement 2.**
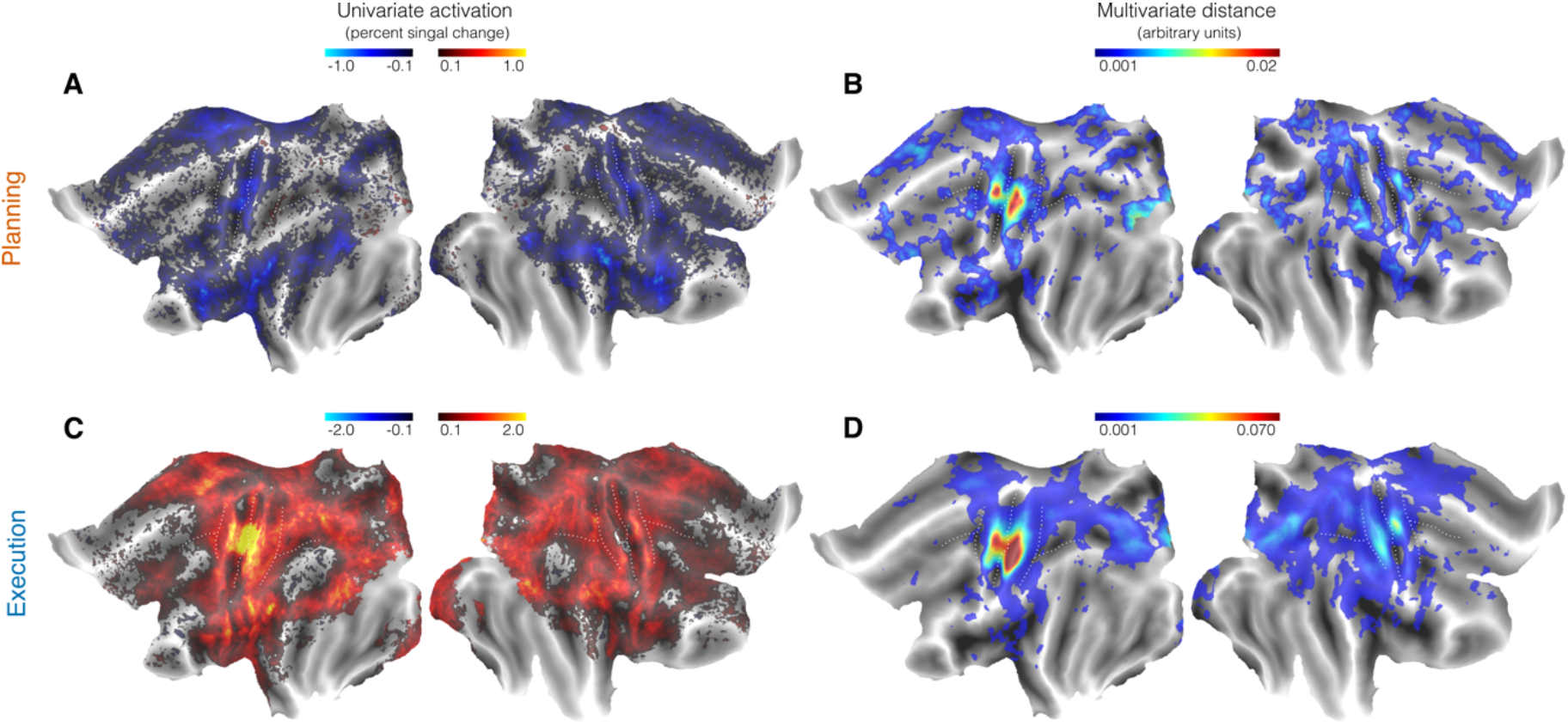
Whole brain flat surface maps (both cortical hemispheres). **A.** Univariate activation for planning (no-go trials). **B.** Multivariate distance for planning (no-go trials). **C.** Univariate activation for execution. **D.** Multivariate distance for execution. Colorbars reflect mean percent signal change for the univariate maps and mean crossnobis distance (arbitrary units) for the multivariate maps.

## References

Alexander GE, Crutcher MD (1990) Neural representations of the target (goal) of visually guided arm movements in three motor areas of the monkey. Journal of Neurophysiology 64:164–178.

Andersen RA, Cui H (2009) Intention, Action Planning, and Decision Making in Parietal-Frontal Circuits. Neuron 63:568–583.

Arbuckle SA, Weiler J, Kirk EA, Rice CL, Schieber M, Pruszynski JA, Ejaz N, Diedrichsen J (2020) Structure of population activity in primary motor cortex for single finger flexion and extension. The Journal of Neuroscience:JN-RM-0999-20.

Ariani G, Diedrichsen J (2019) Sequence learning is driven by improvements in motor planning. Journal of Neurophysiology 121:2088–2100.

Ariani G, Oosterhof NN, Lingnau A (2018) Time-resolved decoding of planned delayed and immediate prehension movements. Cortex 99.

Ariani G, Wurm MF, Lingnau A (2015) Decoding Internally and Externally Driven Movement Plans. Journal of Neuroscience 35:14160–14171.

Azim E, Seki K (2019) Gain control in the sensorimotor system. Current Opinion in Physiology 8:177–187.

Chapman CE, Bushnell MC, Miron D, Duncan GH, Lund JP (1987) Sensory perception during movement in man. Experimental brain research 68:516–524.

Churchland MM, Cunningham JP, Kaufman MT, Ryu SI, Shenoy K V. (2010) Cortical Preparatory Activity: Representation of Movement or First Cog in a Dynamical Machine? Neuron 68:387–400.

Cisek P, Kalaska JF (2004) Neural correlates of mental rehearsal in dorsal premotor cortex. Nature 431:993–996.

Cisek P, Kalaska JF (2010) Neural mechanisms for interacting with a world full of action choices. Annual review of neuroscience 33:269–298.

Coulter JD, Jones EG (1977) Differential distribution of corticospinal projections from individual cytoarchitectonic fields in the monkey. Brain Research 129:335–340.

Crammond DJ, Kalaska JF (2000) Prior information in motor and premotor cortex: activity during the delay period and effect on pre-movement activity. Journal of neurophysiology 84:986–1005.

Cui H, Andersen RA (2007) Posterior Parietal Cortex Encodes Autonomously Selected Motor Plans. Neuron 56:552–559.

Cui H, Andersen RA (2011) Different Representations of Potential and Selected Motor Plans by Distinct Parietal Areas. Journal of Neuroscience 31:18130–18136.

Dale AM, Fischl B, Sereno MI (1999) Cortical Surface-Based Analysis: I. Segmentation and Surface Reconstruction. NeuroImage 9:179–194.

Diedrichsen J, Berlot E, Mur M, Schütt HH, Shahbazi M, Kriegeskorte N (2020) Comparing representational geometries using whitened unbiased-distance-matrix similarity.

Diedrichsen J, Kriegeskorte N (2017) Representational models: A common framework for understanding encoding, pattern-component, and representational-similarity analysis Cichy R, ed. PLOS Computational Biology 13:e1005508.

Diedrichsen J, Yokoi A, Arbuckle SA (2018) Pattern component modeling: A flexible approach for understanding the representational structure of brain activity patterns. NeuroImage 180:119–133.

Ejaz N, Hamada M, Diedrichsen J (2015) Hand use predicts the structure of representations in sensorimotor cortex. Nature Neuroscience 18:1034–1040.

Elsayed GF, Lara AH, Kaufman MT, Churchland MM, Cunningham JP (2016) Reorganization between preparatory and movement population responses in motor cortex. Nature Communications:13239.

Fischl B, Rajendran N, Busa E, Augustinack J, Hinds O, Yeo BTT, Mohlberg H, Amunts K, Zilles K (2008) Cortical folding patterns and predicting cytoarchitecture. Cerebral Cortex 18:1973–1980.

Fischl B, Sereno MI, Tootell RBH, Dale AM (1999) High-resolution intersubject averaging and a coordinate system for the cortical surface. Human Brain Mapping 8:272–284.

Gale DJ, Flanagan JR, Gallivan JP (2021) Human somatosensory cortex is modulated during motor planning. Journal of Neuroscience 41:5909–5922.

Gallivan JP, Johnsrude IS, Flanagan JR (2015) Planning Ahead: Object-Directed Sequential Actions Decoded from Human Frontoparietal and Occipitotemporal Networks. Cerebral Cortex 26:bhu302.

Gallivan JP, McLean DA, Flanagan JR, Culham JC (2013) Where One Hand Meets the Other: Limb-Specific and Action-Dependent Movement Plans Decoded from Preparatory Signals in Single Human Frontoparietal Brain Areas. Journal of Neuroscience 33:1991–2008.

Gallivan JP, McLean DA, Smith FW, Culham JC (2011a) Decoding Effector-Dependent and Effector-Independent Movement Intentions from Human Parieto-Frontal Brain Activity. Journal of Neuroscience 31:17149–17168.

Gallivan JP, McLean DA, Valyear KF, Pettypiece CE, Culham JC (2011b) Decoding Action Intentions from Preparatory Brain Activity in Human Parieto-Frontal Networks. Journal of Neuroscience 31:9599–9610.

Ghez C, Favilla M, Ghilardi MF, Gordon J, Bermejo R, Pullman S (1997) Discrete and continuous planning of hand movements and isometric force trajectories. Experimental Brain Research 115:217–233.

Haith AM, Pakpoor J, Krakauer JW (2016) Independence of Movement Preparation and Movement Initiation. Journal of Neuroscience 36:3007–3015.

Hardwick RM, Forrence AD, Krakauer JW, Haith AM (2019) Time-dependent competition between goal-directed and habitual response preparation. Nature Human Behaviour 3:1252–1262.

Hoshi E, Tanji J (2004) Differential roles of neuronal activity in the supplementary and presupplementary motor areas: from information retrieval to motor planning and execution. Journal of neurophysiology 92:3482–3499.

Hoshi E, Tanji J (2006) Differential involvement of neurons in the dorsal and ventral premotor cortex during processing of visual signals for action planning. Journal of neurophysiology 95:3596–3616.

Hutton C, Bork A, Josephs O, Deichmann R, Ashburner J, Turner R (2002) Image Distortion Correction in fMRI: A Quantitative Evaluation. NeuroImage 16:217–240.

Jafari M, Aflalo T, Chivukula S, Kellis SS, Salas MA, Norman SL, Pejsa K, Liu CY, Andersen RA (2020) The human primary somatosensory cortex encodes imagined movement in the absence of sensory information. Communications biology 3:1–7.

Jiang W, Lamarre Y, Chapman CE (1990) Modulation of cutaneous cortical evoked potentials during isometric and isotonic contractions in the monkey. Brain Research 536:69–78.

Kaufman MT, Churchland MM, Ryu SI, Shenoy K V (2014) Cortical activity in the null space: permitting preparation without movement. Nature Neuroscience 17:440–448.

Keele SW (1968) Movement control in skilled motor performance. Psychological Bulletin 70:387–403.

Keele SW, Summers JJ (1976) The Structure of Motor Programs. In: Motor Control, pp 109–142. Elsevier.

Kikkert S, Pfyffer D, Verling M, Freund P, Wenderoth N (2021) Finger somatotopy is preserved after tetraplegia but deteriorates over time. bioRxiv.

Klapp ST (1995) Motor Response Programming During Simple and Choice Reaction Time: The Role of Practice. Journal of Experimental Psychology: Human Perception and Performance 21:1015–1027.

Klapp ST, Erwin CI (1976) Relation between programming time and duration of the response being programmed. Journal of Experimental Psychology: Human Perception and Performance 2:591–598.

Kuehn E, Mueller K, Turner R, Schütz-Bosbach S (2014) The functional architecture of S1 during touch observation described with 7 T fMRI. Brain Structure and Function 219:119–140.

Ledoit O, Wolf M (2004) Honey, I shrunk the sample covariance matrix. The Journal of Portfolio Management 30:110–119.

Leoné FTM, Heed T, Toni I, Medendorp WP (2014) Understanding effector selectivity in human posterior parietal cortex by combining information patterns and activation measures. The Journal of neuroscience 34:7102–7112.

Logothetis NK, Pauls J, Augath M, Trinath T, Oeltermann A (2001) Neurophysiological investigation of the basis of the fMRI signal. Nature 412:150–157.

Oosterhof NN, Wiestler T, Downing PE, Diedrichsen J (2011) A comparison of volume-based and surface-based multi-voxel pattern analysis. NeuroImage 56:593–600.

Oosterhof NN, Wiggett AJ, Diedrichsen J, Tipper SP, Downing PE, Ramsey R, Hamilton AFDC (2012) Surface-Based Information Mapping Reveals Crossmodal Vision – Action Representations in Human Parietal and Occipitotemporal Cortex Surface-Based Information Mapping Reveals Crossmodal Vision - Action Representations in Human Parietal and Occipitotemporal. Journal of neurophysiology 104:1077–1089.

Puckett AM, Bollmann S, Barth M, Cunnington R (2017) Measuring the effects of attention to individual fingertips in somatosensory cortex using ultra-high field (7T) fMRI. NeuroImage 161:179–187.

Rathelot J-A, Strick PL (2006) Muscle representation in the macaque motor cortex: An anatomical perspective. Proceedings of the National Academy of Sciences 103:8257 LP–8262.

Riehle A, Requin J (1989) Monkey primary motor and premotor cortex: single-cell activity related to prior information about direction and extent of an intended movement. Journal of neurophysiology 61:534–549.

Rosenbaum DA (1980) Human movement initiation: Specification of arm, direction, and extent. Journal of Experimental Psychology: General 109:444–474.

Seki K, Fetz EE (2012) Gating of sensory input at spinal and cortical levels during preparation and execution of voluntary movement. Journal of Neuroscience 32:890–902.

Shenoy K V, Sahani M, Churchland MM (2013) Cortical control of arm movements: a dynamical systems perspective. Annual Review of Neuroscience 36:337–359.

Starr A, Cohen LG (1985) ‘Gating’ of somatosensory evoked potentials begins before the onset of voluntary movement in man. Brain Research 348:183–186.

Tanji J, Evarts E V. (1976) Anticipatory activity of motor cortex neurons in relation to direction of an intended movement. Journal of Neurophysiology 39:1062–1068.

Turella L, Tucciarelli R, Oosterhof NN, Weisz N, Rumiati R, Lingnau A (2016) Beta band modulations underlie action representations for movement planning. NeuroImage 136:197–207.

Umeda T, Isa T, Nishimura Y (2019) The somatosensory cortex receives information about motor output. Science Advances 5.

Vyas S, Golub MD, Sussillo D, Shenoy K V (2020) Computation Through Neural Population Dynamics. Annual Review of Neuroscience 43:249–275.

Walther A, Nili H, Ejaz N, Alink A, Kriegeskorte N, Diedrichsen J (2016) Reliability of dissimilarity measures for multi-voxel pattern analysis. NeuroImage 137:188–200.

Widener GL, Cheney PD (1997) Effects on Muscle Activity From Microstimuli Applied to Somatosensory and Motor Cortex During Voluntary Movement in the Monkey. Journal of Neurophysiology 77:2446–2465.

Wong AL, Haith AM (2017) Motor planning flexibly optimizes performance under uncertainty about task goals. Nature Communications 8:14624.

Yokoi A, Arbuckle SA, Diedrichsen J (2018) The role of human primary motor cortex in the production of skilled finger sequences. Journal of Neuroscience 38:1430–1442.

Yousry T (1997) Localization of the motor hand area to a knob on the precentral gyrus. A new landmark. Brain 120:141–157.

