## Supplementary figures and images for "Motor planning brings human primary somatosensory cortex into action-specific preparatory states"

### Supplemental Data 1

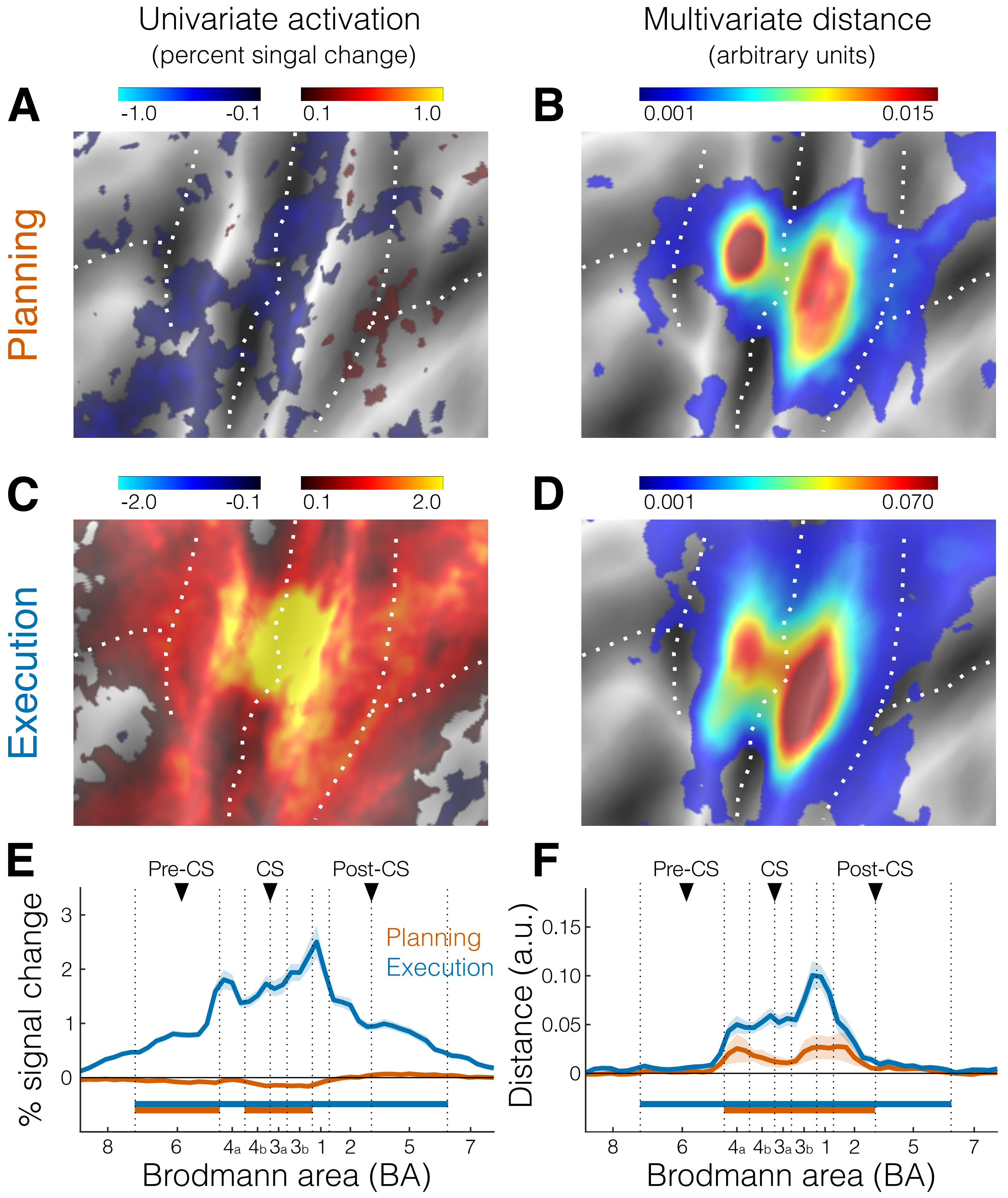

### Supplemental Data 2

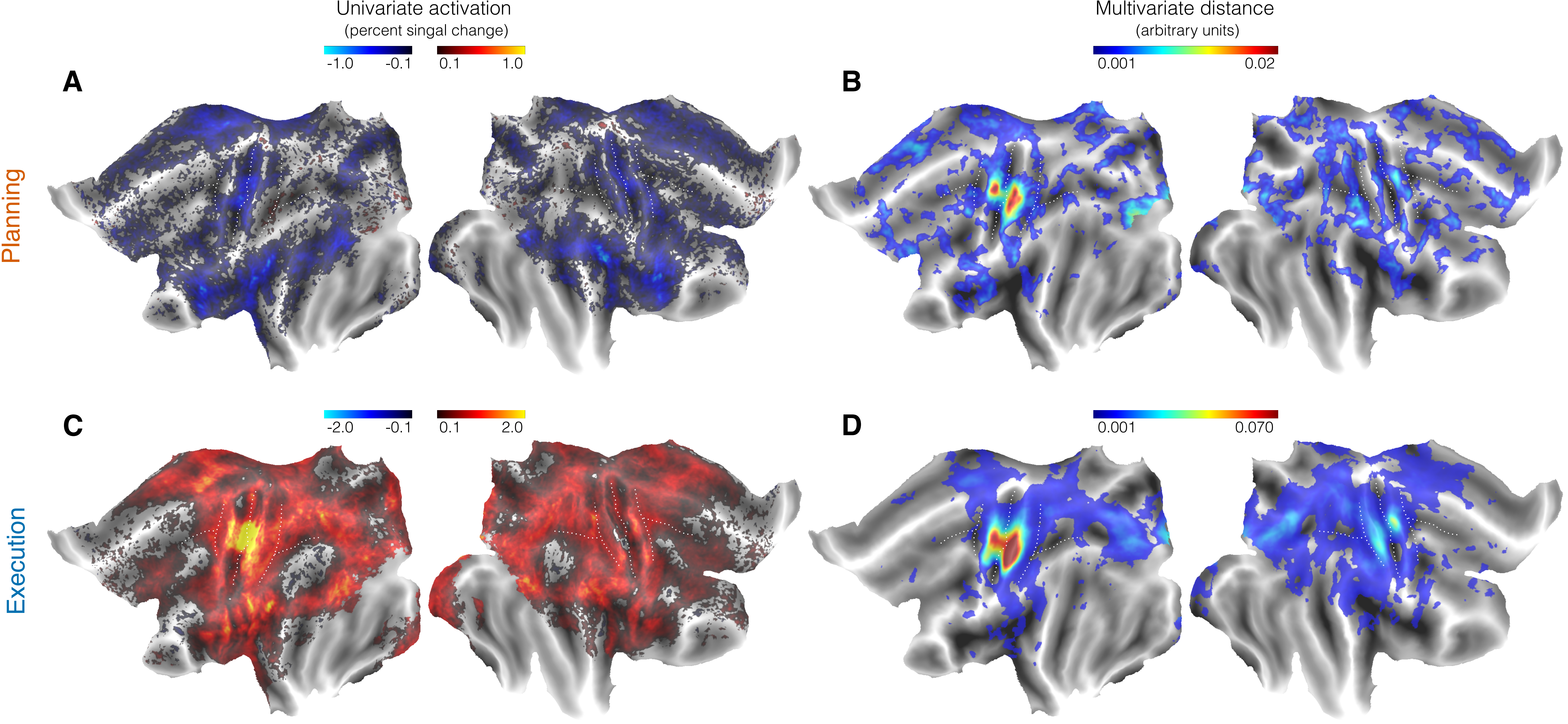
